# Liver ductal organoids reconstruct intrahepatic biliary trees in decellularized liver grafts

**DOI:** 10.1101/2021.11.14.468338

**Authors:** Katsuhiro Tomofuji, Ken Fukumitsu, Jumpei Kondo, Hiroshi Horie, Kenta Makino, Satoshi Wakama, Takashi Ito, Yu Oshima, Satoshi Ogiso, Takamichi Ishii, Masahiro Inoue, Etsuro Hatano

**Affiliations:** Department of Clinical Bio-resource Research and Development, Graduate School of Medicine, Kyoto University, 46-29, Shimoadachi-cho, Sakyo-ku, Kyoto, 606-8501, Japan; Department of Surgery, Graduate School of Medicine, Kyoto University, 54 Kawahara-cho, Shogoin, Sakyo-ku, Kyoto, 606-8507, Japan

**Author notes:** Corresponding author. Department of Clinical Bio-resource Research and Development, Graduate School of Medicine, Kyoto University, 46-29, Shimoadachi-cho, Sakyo-ku, Kyoto, 606-8501, Japan, *E-mail address. Corresponding author. Department of Surgery, Graduate School of Medicine, Kyoto University, 54 Kawahara-cho, Shogoin, Sakyo-ku, Kyoto, 606-8507, Japan, *E-mail address. J.K. Division of Health Sciences, Department of Molecular Biology and Clinical Investigation, Graduate School of Medicine, Osaka University, Yamadaoka, Suita City, Osaka 565-0871, Japan.

**Keywords:** Decellularization, Recellularization, Bile ducts, Tissue engineering, Extracellular matrix, Organoids

## Abstract

Three-dimensional scaffolds decellularized from native organs are a promising technique to establish engineered liver grafts and overcome the current shortage of donor organs. However, limited sources of bile duct cells and inappropriate cell distribution in bioengineered liver grafts have hindered their practical application. Organoid technology is anticipated to be an excellent tool for the advancement of regenerative medicine. In the present study, we reconstructed intrahepatic bile ducts in a rat decellularized liver graft by recellularization with liver ductal organoids. Using an *ex vivo* perfusion culture system, we demonstrated the biliary characteristics of repopulated mouse liver organoids, which maintained bile duct markers and reconstructed biliary tree-like networks with luminal structures. We also established a method for the co-recellularization with engineered bile ducts and primary hepatocytes, revealing the appropriate cell distribution to mimic the native liver. We then utilized this model in human organoids to demonstrate the reconstructed bile ducts. Our results show that liver ductal organoids are a potential cell source for bile ducts from bioengineered liver grafts using three-dimensional scaffolds.

## 1. Introduction

Liver transplantation is currently the only curative option for patients with end-stage liver disease. However, the demand for liver organs greatly exceeds the supply of donor livers. To address this challenge, approaches such as cell transplantation, bioartificial organs, and liver support devices have been explored [1, 2]; however, none have yet been established as therapeutic alternatives.

Decellularization and recellularization, in which an extracellular matrix (ECM) is prepared from its native organs, retaining the inherent structure and biological properties, followed by recellularization with new cells to create transplantable functional organs, are promising techniques for tissue engineering [3]. Since the first report of a decellularized liver scaffold [4], recellularized liver graft models have been investigated from some liver cell sources. In addition to hepatocytes, a major functional cell type to be recellularized, the source cells include cholangiocytes and endothelial cells, as well as other stromal cells. Hepatocyte cell sources have been widely reported, and include primary hepatocytes, fetal hepatocytes, induced pluripotent stem cells (iPSC)-derived hepatocyte-like cells, and direct reprogramming of fibroblasts [5-9]. In contrast, the study of cholangiocyte cell sources has been rare owing to the difficulty of culturing primary cholangiocytes. Mouse immortalized cholangiocytes [10], normal rat cholangiocytes [11], and iPSC-derived cholangiocytes [9] have been studied as sources to recellularize the decellularized liver tissue ECM. Although there is proof-of-concept to show that external cells can repopulate decellularized liver tissue, further physiological cholangiocyte candidates are needed for cellular sources.

Recent advances have enabled the culture of tissue stem cells as three-dimensional (3D) organoids, which self-organize into 3D structures mimicking the original organs [12]. This organoid culture technique has also been applied to liver stem cells. A bile duct fragment embedded in Matrigel self-organized into liver ductal organoids with bipotential differentiation capacity into both hepatocyte and cholangiocyte lineages [13, 14]. Liver ductal organoids have been reported as a potential cell source for hepatocyte regeneration [15]. Therefore, we expect that the liver ductal organoids may also be a cell source for the regeneration of cholangiocytes, and the proliferating and self-organizing ability of the organoids provide a means to obtain cell networks for tissue engineering.

Here, we have described liver ductal organoids as a potential bile duct cell source for a bioengineered liver graft. We characterized the biliary properties of liver ductal organoids *in vitro* and those of repopulated bile ducts in a bioengineering liver graft *ex vivo*. The appropriate cell placement and structure of the recellularized liver also demonstrated advantages for future clinical progress.

## 2. Results

### 2.1. Mouse liver ductal organoids exhibit characteristics of functional cholangiocytes

Although liver ductal organoids are derived from intrahepatic bile ducts, these cells have mainly been investigated as a resource for differentiated hepatocytes [13, 14, 16-18] rather than for cholangiocytes [19, 20]. We investigated to what extent the ductal organoids are characteristic of cholangiocytes. Liver ductal organoids showed rapid proliferation when cultured in Matrigel (Fig. 1A–C). Liver ductal organoids under maintenance culture were analyzed by RT-qPCR and immunofluorescence to determine the expression of markers specific to cholangiocytes and hepatocytes. In the RT-qPCR analyses, the expression of the cholangiocyte-specific markers *Krt19* and *Sox9* was 0.68-fold lower and 2.0-fold higher in ductal organoids than extrahepatic bile ducts, respectively (Fig 1D). In contrast, the expression of hepatocyte-specific markers, *Alb, Hnf4a*, and *Cyp3a11*, was quite low compared with that in primary hepatocytes. Immunofluorescence analyses revealed that ductal organoids exhibit cystic structures expressing cholangiocyte lineage markers, such as KRT19, SOX9, ASBT, and CFTR, whereas ALB was absent (Fig. 1E). These findings indicate that liver ductal organoids under maintenance culture conditions, which reportedly possess bipotent stemness, exhibit characteristics of cholangiocytes but not of hepatocytes.

**Fig. 1.**
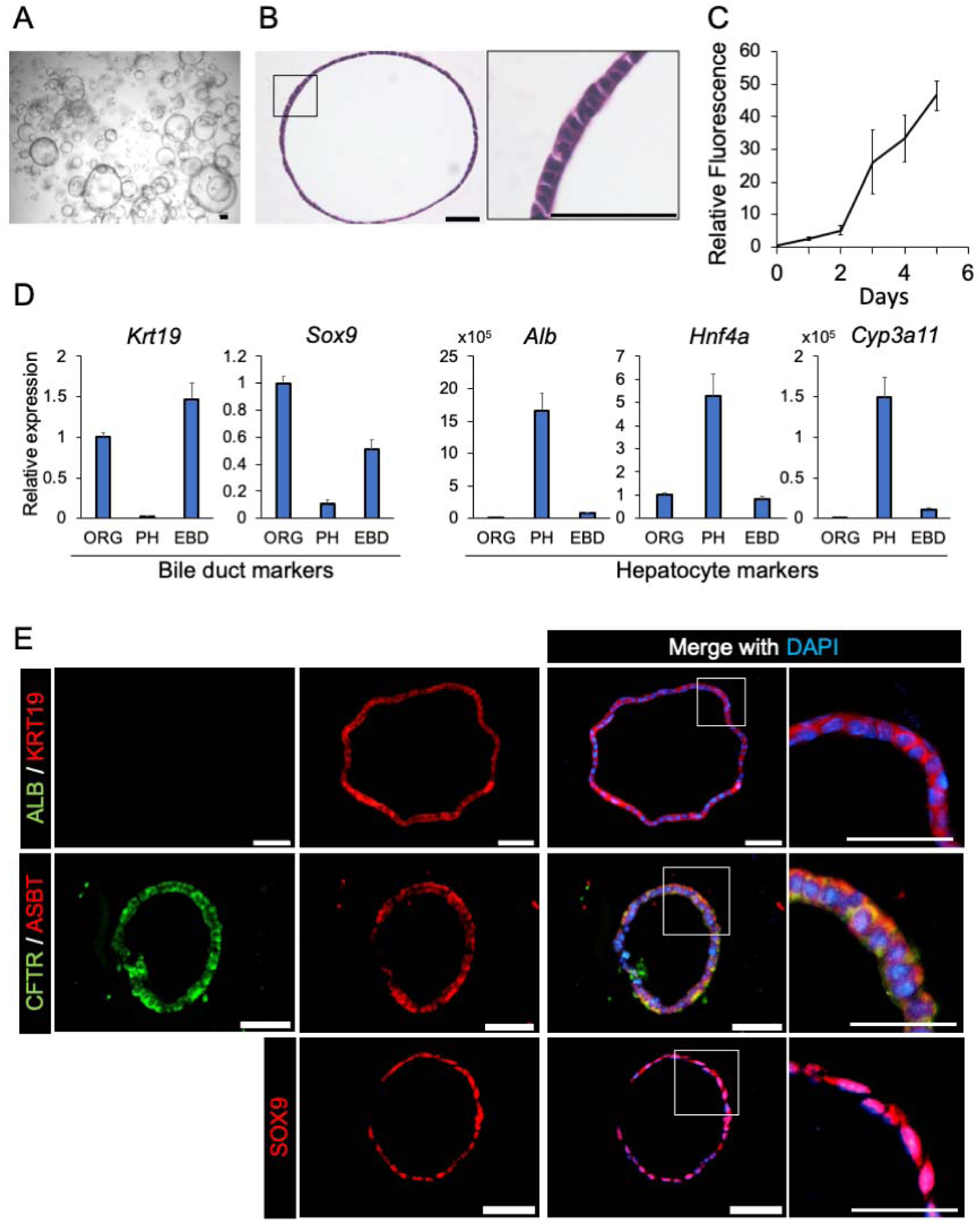
Characterization of mouse liver ductal organoids. (A) Bright-field image of mouse liver ductal organoids cultured in Matrigel. Scale bars: 100 μm. (B) Images of hematoxylin and eosin (HE) staining. A higher magnification image of the outlined box is shown in the right panel. Scale bars: 50 μm. (C) Time course of the viability of the cultured ductal organoids evaluated by ATP assay. The values are the average ± SD. N = 3 for each condition. (D) Relative gene expression analyzed by RT-qPCR in the liver ductal organoids (ORG), primary hepatocytes (PH), and extrahepatic bile duct tissue (EBD). The values are the average ± SD. N = 3 for each condition. (E) Immunofluorescence staining of ductal organoids labeled with the indicated cholangiocyte and hepatocyte markers. Higher magnification images of the white outlined boxes are shown in the right panels. Scale bars: 50 μm.

We subsequently characterized liver ductal organoids by focusing on their cholangiocyte function. The multidrug resistance protein 1 (MDR1) transporter is expressed in normal biliary epithelia and is involved in the efflux of a broad ranges of substrates into the lumen [21]. We thus evaluated the ability of organoids to efflux rhodamine 123, which is mainly transported by MDR1 transporters. Liver ductal organoids were incubated with the dye rhodamine 123 (Fig. 2A), and the dye was transported into the lumen of the cystic organoids. In contrast, in the presence of 20 μM verapamil, which inhibits MDR1 function, rhodamine 123 did not accumulate in the lumen (Fig. 2A). Quantification of the fluorescence intensity showed that verapamil significantly blocked the transportation of rhodamine 123 (*P* value < 0.0001) (Fig. 2B), indicating that this fluorescent dye was actively transported by MDR1.

**Fig. 2.**
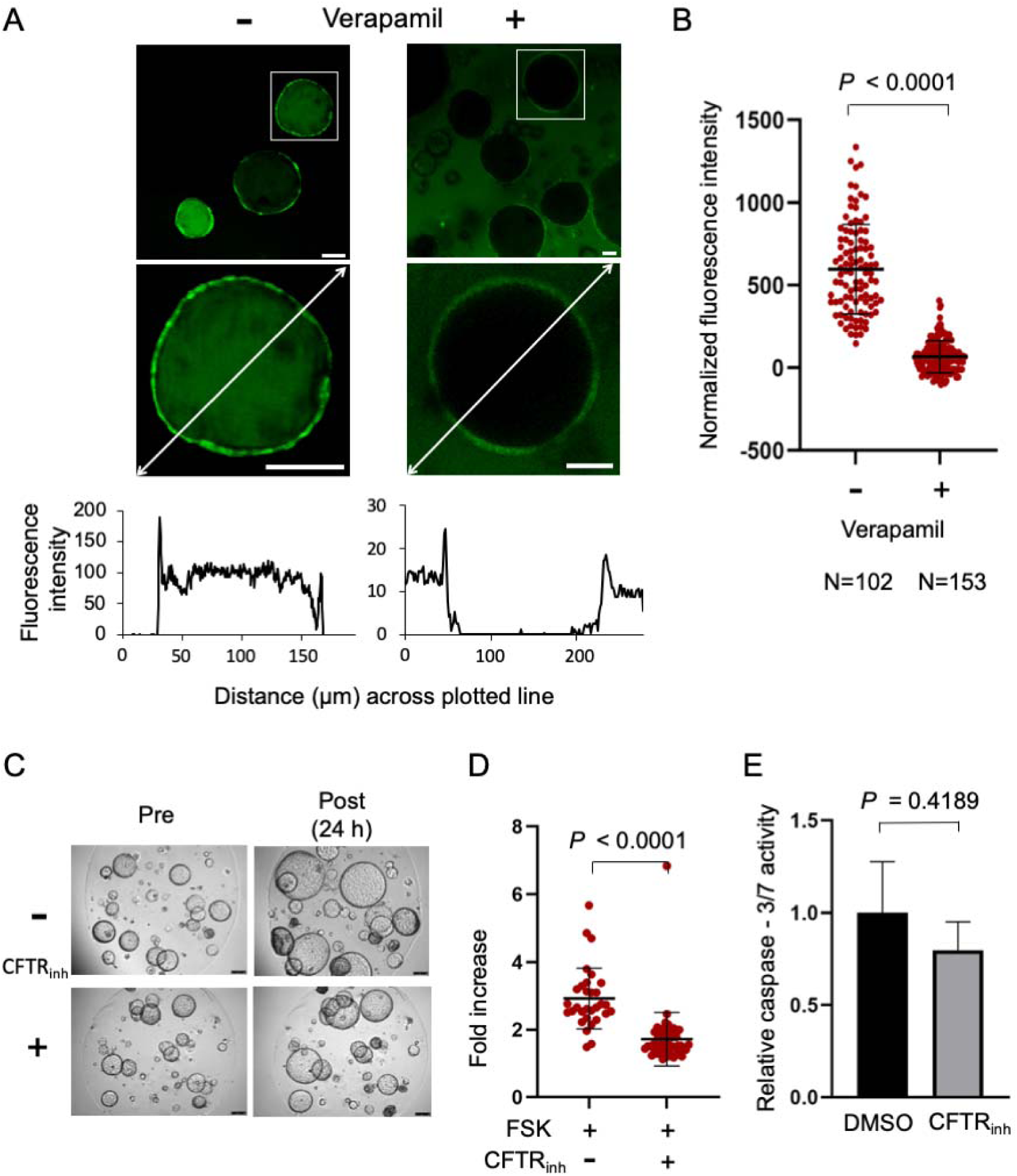
Mouse liver ductal organoids exhibit functional characteristics of cholangiocytes. (A) Representative images by confocal microscopy showing uptake of rhodamine 123 dye in liver ductal organoids in the absence (left) or presence (right) of verapamil, an MDR1 inhibitor. Higher magnification images of the outlined boxed are shown below. Graphs below the fluorescence images depict the fluorescence intensity along the white line in each image. Scale bars: 50 μm. (B) Quantification of mean intraluminal fluorescence intensity normalized to background levels for the organoids in the absence or presence of verapamil over the first 30 minutes. (C) Representative bright-field images in the absence or presence of CFTRinh-172, a CFTR inhibitor, before treatment and after incubation for 24 h. Scale bars: 250 μm. (D) Quantification of the cyst swelling rate after forskolin (FSK) stimulation in the absence or presence of CFTRinh-172, a CFTR inhibitor. N=31 and N=51 for control and CFTRinh-172–treated groups, respectively. (E) Comparison of caspase activation after treatment with DMSO or CFTRinh-172. The values are the average ± SD. N = 3 for each condition.

In addition, to assess the function of another transporter expressed in normal biliary epithelia, cystic fibrosis transmembrane conductance regulator (CFTR), we conducted a forskolin-induced swelling assay. This assay evaluates the increase in CFTR function induced by the activation of cAMP pathway, which is stimulated by forskolin, and leads to cyst swelling [22, 23]. Liver ductal organoids were incubated with forskolin in the absence or presence of CFTR_inh_-172, a specific CFTR inhibitor (Fig. 2C). Organoids increased the cyst size by 2.66-fold after forskolin treatment (Fig. 2D). This forskolin-induced swelling ratio was significantly reduced to 1.55 by CFTR_inh_-172 (*P* value < 0.0001), indicating that liver ductal organoids have CFTR functional activity. The inhibition of CFTR was not associated with cell death (*P* value > 0.4) (Fig. 2E). These findings suggest that liver ductal organoids have the properties of functional cholangiocytes.

### 2.2. Mouse liver ductal organoid cells repopulate the decellularized rat liver ECM to reconstruct biliary trees

As the ductal organoids exhibit the characteristics of cholangiocytes to certain extent, we then investigated if the ductal organoids were a potential cell source for bioengineered livers. We have previously described a decellularized liver graft technique that offers a bioengineered scaffold with a physiological ECM [6, 8] (Supplementary Fig. S1). In addition to the native ECM, decellularized liver also retains the vascular and biliary network frame structure (Supplementary Fig. S2). Histological analysis confirmed the absence of cells in the decellularized liver scaffold (Supplementary Fig. S3). After confirming that the biliary structure was preserved in the decellularized liver, liver ductal organoid cells were then injected into the biliary network of a decellularized rat whole liver scaffold via the common bile duct (Fig. 3A). The recellularized scaffold was cultured in an *ex vivo* perfusion culture system for 3–5 days (Fig. 3B); then, the tissue was formalin-fixed and paraffin-embedded for histological analysis. The ductal organoid-derived cells engrafted along the bile duct walls, forming a monolayered structure lining the lumens (Fig. 3C). RT-qPCR analyses revealed that recellularized bile ducts expressed cholangiocyte marker genes, including *Krt19, Sox9, Cftr*, and *Hnf1*β at comparable levels with those in the extrahepatic bile duct *in vivo* (Fig. 3D), whereas hepatocyte markers (*Alb, Cyp3a11, Hnf4a*) remained at a low level. Moreover, the repopulated organoid cells sustained the expression of stemness markers (*Lgr5, Prom1*). In the immunofluorescence analyses, the repopulated organoid cells also exhibited key biliary markers (KRT19, SOX9, ASBT, CFTR) (Fig. 4A). PCNA, a proliferation marker originally positive in ductal organoids, was maintained in some areas even after recellularization (Fig. 4B), indicating that the engrafted cells retained proliferative ability in the bile duct structure. From these findings, it was determined that repopulated liver ductal organoids were engrafted along the bile duct ECM, maintaining cholangiocyte properties.

**Fig. 3.**
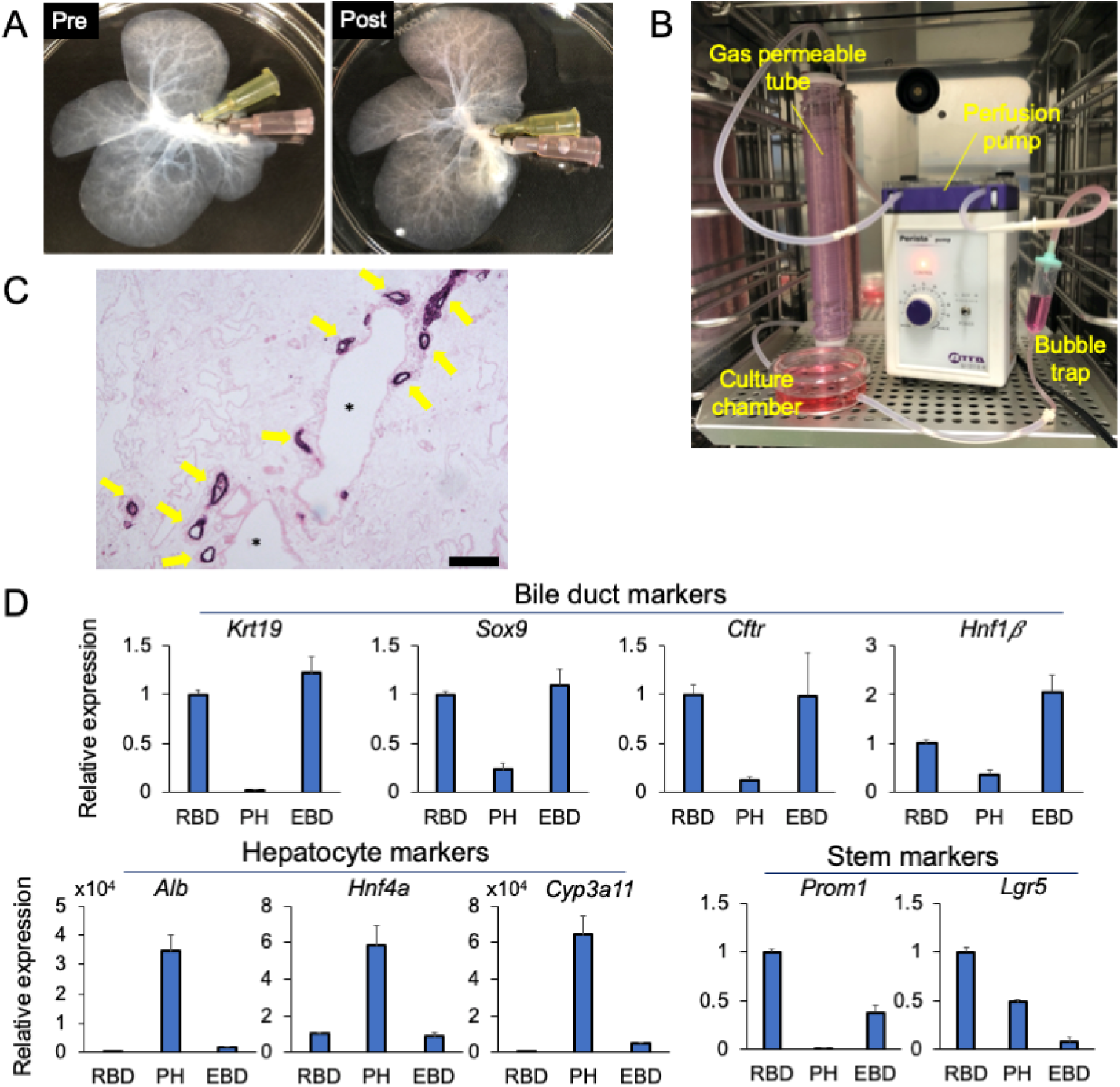
Characterization of recellularized bile ducts. (A) Macroscopic images of a recellularized liver graft before (left) and after (right) recellularization in the bile ducts. (B) Photograph of the perfusion culture set-up placed in a CO_2_ incubator. (C) HE staining of the recellularized bile ducts. Scale bars: 200 μm. (D) Relative gene expression, as analyzed by RT-qPCR, in the recellularized bile ducts (RBD), primary hepatocytes (PH), and extrahepatic bile duct tissue (EBD). The values are the average ± SD. N = 3 for each condition.

**Fig. 4.**
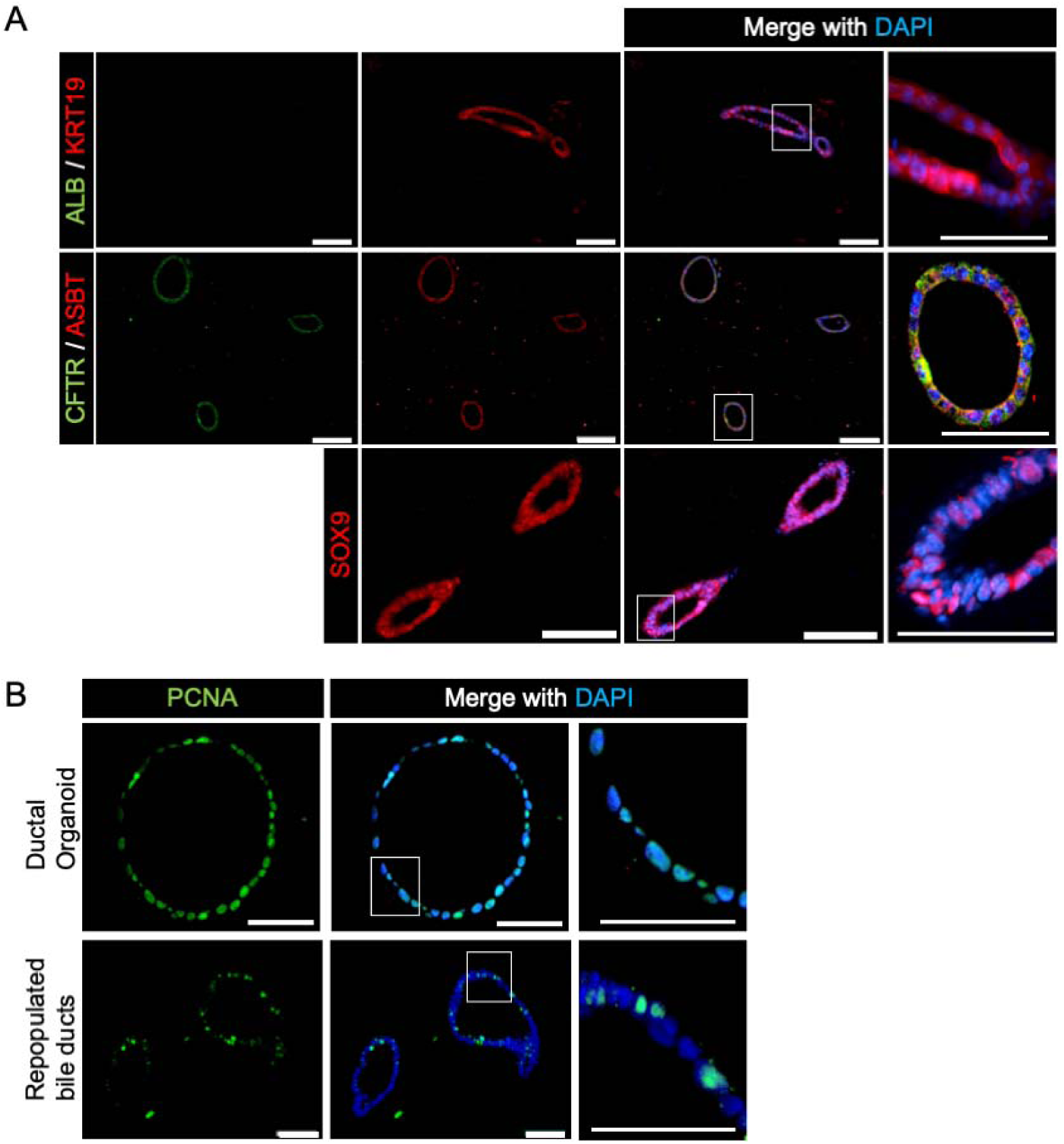
The recellularized bile ducts maintain biliary properties and proliferation ability. (A) Immunofluorescence staining images of the recellularized bile ducts labeled with the indicated cholangiocyte and hepatocyte markers. Scale bars: 100 μm. Higher magnification images of the white outlined boxes are shown in the right panels. Scale bars in high-magnification images: 50 μm. (B) Immunofluorescence images of the recellularized bile ducts (upper panels) and liver ductal organoids (lower panels) stained with PCNA. Higher magnification images of the white outlined boxes are shown in the right panels. Scale bars: 50 μm.

Next, to demonstrate the 3D structure of recellularized bile ducts, ductal organoid cells expressing GFP were inoculated into a decellularized liver. The cells spread to the periphery of the liver, displaying a branched tree-like network (Fig. 5A). In addition, the confocal microscopy images showed engrafted cells on the decellularized bile duct ECM that formed luminal structures (Fig. 5B and Supplementary video S1). As shown by these results, the intrahepatic bile ducts were recellularized efficiently by the injection of ductal organoid-derived cells via the common bile duct.

**Fig. 5.**
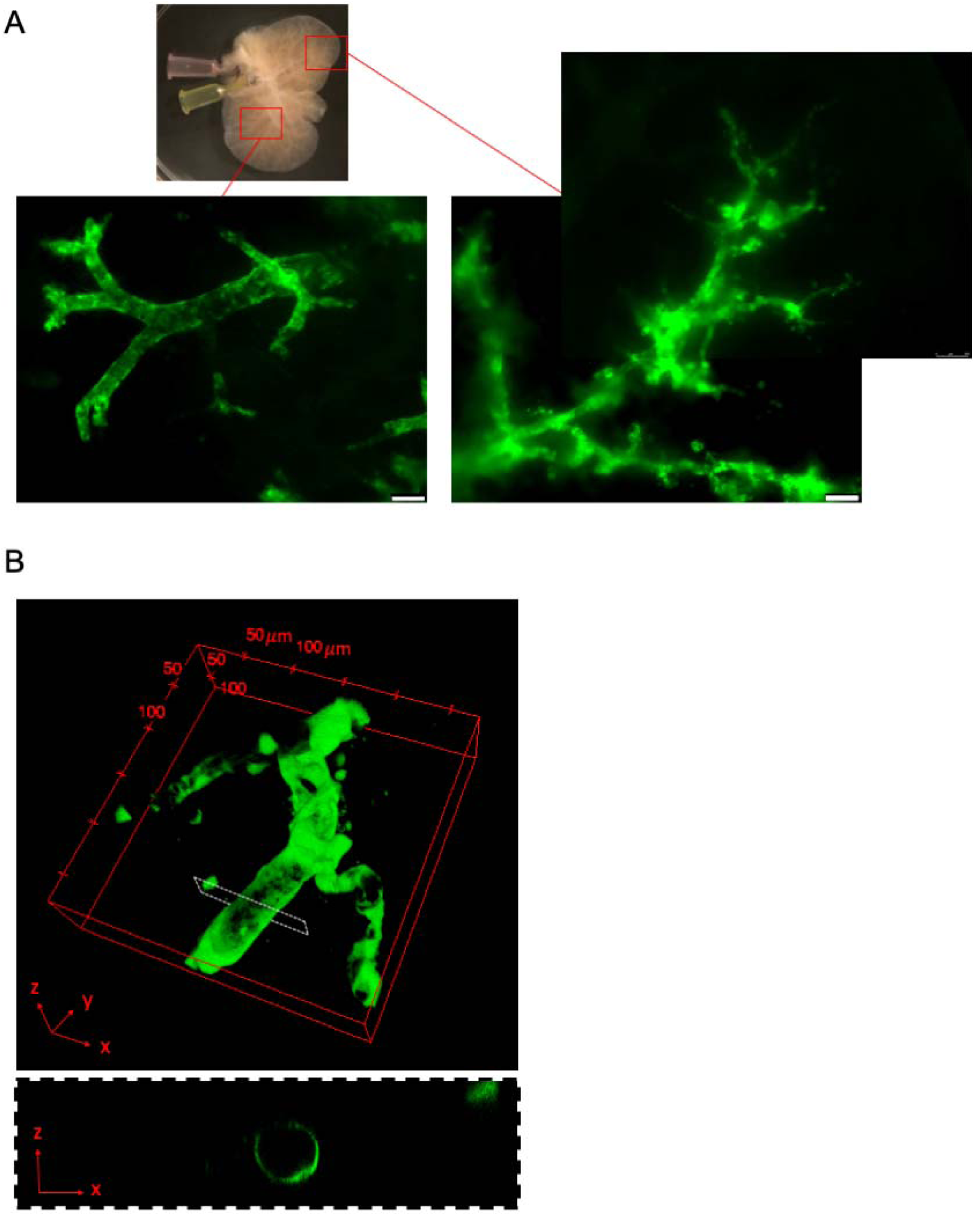
Recellularized bile ducts reconstruct biliary tree-like structures. (A) Fluorescent microscopic images of the bile ducts recellularized with liver ductal organoids expressing GFP on day 5 of perfusion culture. Scale bars: 250 μm. (B) Upper panel: 3D reconstruction of laser confocal microscopy images of the bile ducts recellularized with liver ductal organoids expressing GFP. Lower panel: cross-sectional view (xz) of the white dashed box in the upper panel.

### 2.3. Liver ductal organoid cells retain cholangiocyte characteristics during simultaneous recellularization with primary hepatocytes

Hepatocytes are the main functional cellular unit in the liver. We evaluated whether the co-recellularization of repopulated liver ductal organoids and hepatocytes affected their engraftment or differentiation properties. As the optimal culture medium for hepatocytes and cholangiocytes differs, we first injected 5 × 10^6^ liver ductal organoid-derived cells via the common bile duct and the recellularized liver was perfused with expansion medium (EM) containing 10 μM forskolin (Supplementary Table S1), a bile duct organoid EM, via the portal vein for 5 days. Then, freshly isolated mouse primary hepatocytes (5 × 10^7^ cells) were injected via the common bile duct, followed by perfusion culture with HCM^™^ (Lonza Sales Ltd, Basel, Switzerland), a hepatocyte culture medium, via the portal vein for 2 days (Fig. 6A). Histological analyses of this co-recellularized liver revealed the appropriate cell distribution of hepatocytes into the parenchymal space and of liver ductal organoid cells into the bile duct (Fig. 6B, C). Through the immunofluorescence analyses, repopulated primary hepatocytes were shown to express ALB and HNF4a. Repopulated liver ductal organoid cells expressed KRT19 and SOX9 but not ALB or HNF4a (Fig. 6D), showing that the repopulated cells maintained biliary lineage and did not differentiate to hepatocyte lineage when co-cultured with primary hepatocytes.

**Fig. 6.**
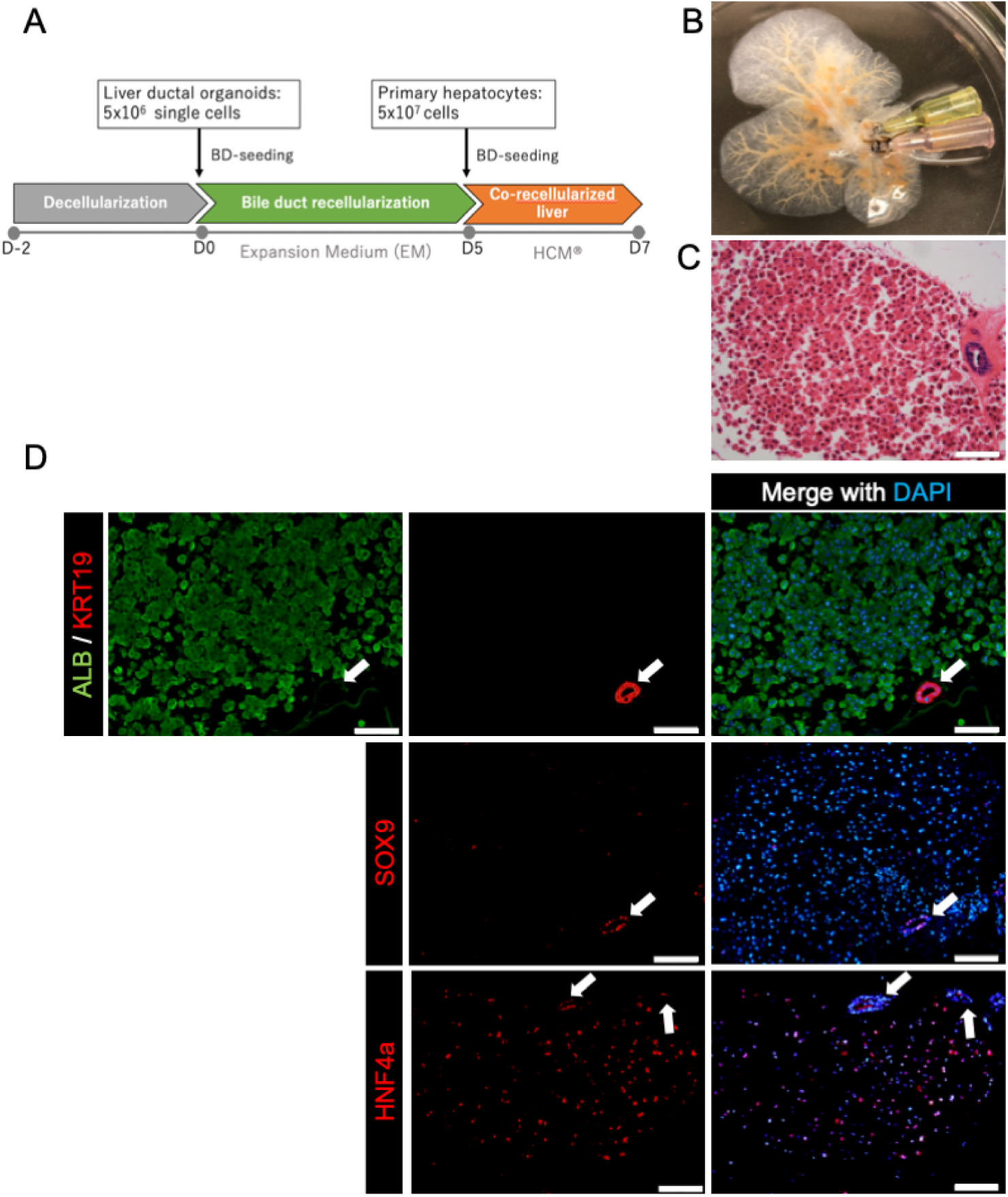
Co-recellularization of engineered bile ducts and primary hepatocytes results in distinguished localization and maintenance of each specific cell markers. (A) The scheme for co-recellularization of mouse liver ductal organoids and mouse primary hepatocytes. BD-seeding: Cells were injected via the common bile duct. (B) Macroscopic images of the co-recellularized liver with primary hepatocytes and ductal organoids. (C) HE staining of the co-recellularized liver on day 7. Scale bar: 100 μm. (D) Immunofluorescence staining of the indicated cholangiocyte and hepatocyte markers. White arrows indicate recellularized bile ducts. Scale bars: 100 μm.

### 2.4. Human liver ductal organoid cells are capable of repopulating decellularized rat liver ECM

These findings must be extended to human cells to establish a transplantable human liver graft in the future. Human liver ductal organoids have bipotential capacity to differentiate into both hepatocytes and cholangiocytes, similar to mouse liver ductal organoids [14]. We generated three liver ductal organoid lines from residual liver specimens from patients undergoing hepatectomy in response to liver tumors (Table 1). Human liver ductal organoids cultured in Matrigel (Fig. 7A, B) expressed key biliary markers (KRT19, SOX9) but not ALB (Fig. 7C). Next, we injected the human organoid cells into rat decellularized liver and achieved successful recellularization (Fig. 7D), similar to the mouse organoids, in a rat decellularized liver. The RT-qPCR analysis demonstrated that *KRT19* and *PROM1* were upregulated in all cases evaluated and *SOX9* was upregulated in two of three cases (Fig. 7E). In contrast, the gene expression of hepatocyte markers (*ALB, CYP3A4*) remained low (Fig. 7E). In the immunofluorescence analyses, the recellularized bile ducts were shown to maintain the expression of KRT19 and SOX9 (Fig. 7F). Therefore, human liver ductal organoids maintained cholangiocyte properties after recellularization.

**Table 1.** Patient data for organoids and tissue

| No. | I.D. | Age | Sex | Background disease |
| --- | --- | --- | --- | --- |
| 1 | KUH2N | 59 | M | Colorectal liver metastases |
| 2 | KUH6N | 71 | M | Intrahepatic cholangiocarcinoma |
| 3 | KUH7N | 16 | F | Focal nodular hyperplasia |

**Fig. 7.**
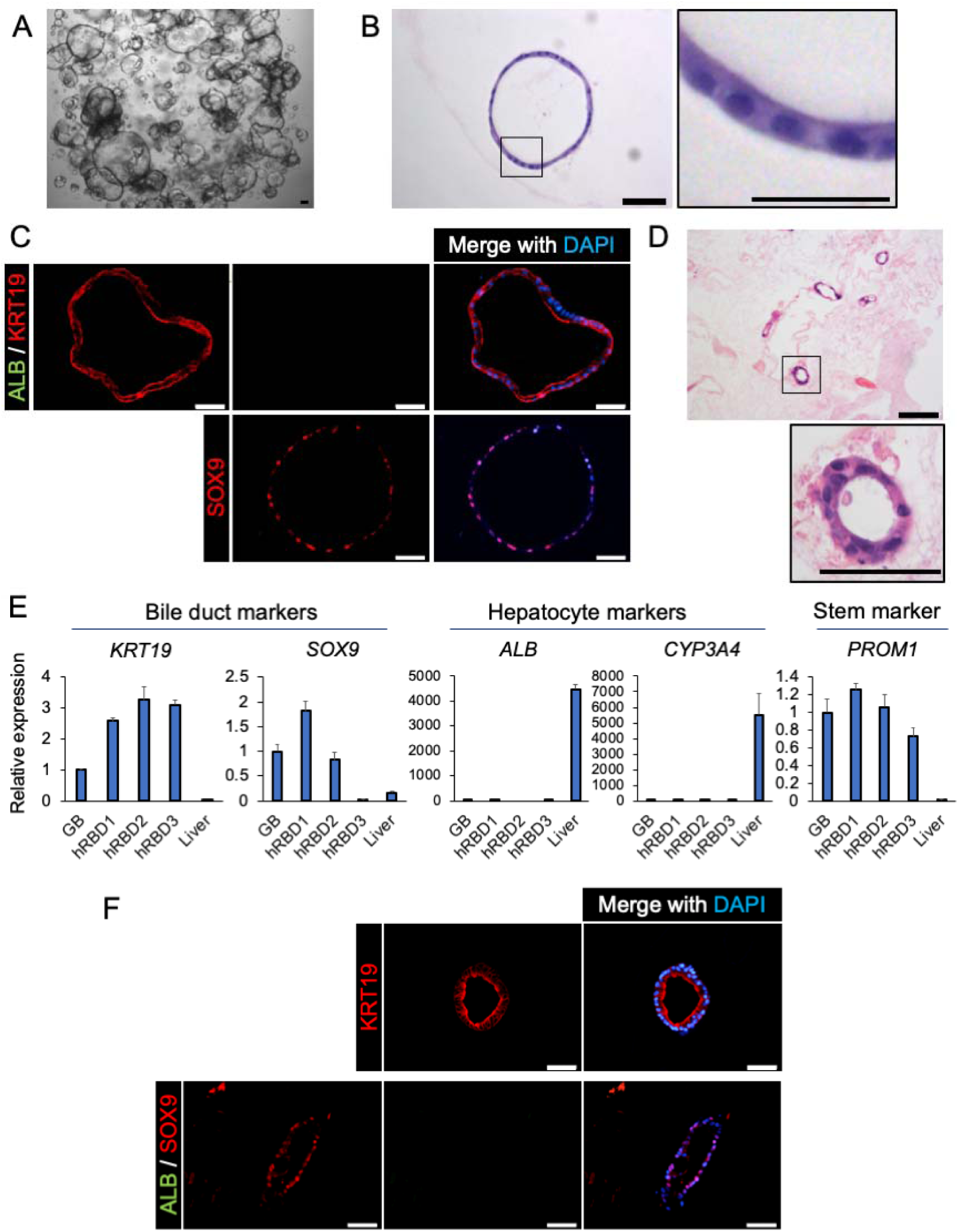
Human liver ductal organoids can repopulate decellularized bile ducts. (A) Bright-field image of human liver ductal organoids cultured in Matrigel. Scale bars: 100 μm. (B) HE staining. Higher magnification images of the outlined boxes are shown in the right panel. Scale bars: 50 μm (left), 25 μm (right). (C) Immunofluorescence staining of ductal organoids cultured in Matrigel labeled with the indicated cholangiocyte and hepatocyte markers. Scale bars: 50 μm. (D) A HE staining image of bile ducts recellularized with human liver ductal organoids. Higher magnification images of the outlined boxes are shown in the lower panel. Scale bars: 100 μm (upper), 50 μm (lower). (E) Relative gene expression analyzed by RT-qPCR of gallbladder cholangiocytes, recellularized bile ducts of each human organoid line (hRBD1, KUH2N; hRBD2, KUH6N; hRBD3, KUH7N), and liver tissue. The values are the average ± SD. N = 3 for each condition. (F) Immunofluorescence staining of bile ducts recellularized with human liver ductal organoids and labeled with the indicated cholangiocyte and hepatocyte markers. Scale bars: 50 μm.

## 3. Discussion

Here, we have reported that liver ductal organoids can be a useful cholangiocyte cell source for bioengineered bile ducts. Luminal structures with a single cell layer lining were successfully reconstituted in the 3D scaffold decellularized from the liver which expressed appropriate bile duct markers. In addition, the co-recellularization with both primary hepatocytes and liver ductal organoids enabled the proper cell placement similar to the native liver, maintaining specific markers for each cell type.

Since the liver ductal organoid was first introduced as a bipotential stem cell culture [13], hepatocyte lineage differentiation from the ductal organoids has been well studied [17, 18]. Very recently, several papers investigating the cholangiocyte characteristics of this bipotent organoid were published [19, 20, 24]. The induction of biliary lineage differentiation in liver ductal organoids upregulated cholangiocyte-related genes and downregulated stem cell markers such as *Lgr5* and *Ki67* [19]. However, in this study, we found that the liver ductal organoids already possess cholangiocyte-like properties prior to differentiation, including representative cholangiocyte markers and functions. In addition, the repopulated liver ductal organoid-derived cells remained proliferative and retained the expression of stem cell markers *Lgr5* and *Prom1* as well as key biliary markers (*Krt19* and *Sox9)*. This characteristic appears to be a favorable feature as a source for tissue engineering because maintaining the proliferating potential is important to enable engrafted cells to expand and self-organize in a decellularized liver graft *ex vivo*, or even after transplantation. Liver ductal organoids morphologically consisted of flat cells forming a cystic structure in vitro, meanwhile the recellularized bile ducts showed luminal structures in the form of a simple columnar epithelium. We also confirmed that repopulated liver ductal organoids were not differentiated into hepatocyte lineage, despite their bipotential capacity and perfusion with hepatocyte medium in the co-recellularized liver.

With regard to cholangiocyte cell sources for recellularization, a similar approach was recently reported [20], in which engineered extrahepatic bile duct models *in vitro* were established using human decellularized extrahepatic bile ducts repopulated with bile duct-derived organoids. They concluded that among three types of bile duct-derived organoids, namely liver ductal organoids, extrahepatic bile duct-derived organoids, and bile-derived organoids, extrahepatic bile duct-derived and bile-derived organoids repopulated decellularized extrahepatic bile ducts efficiently *in vitro*. While the liver ductal organoid was a promising extrahepatic bile duct cell source, our study demonstrated that liver ductal organoids from both mice and humans successfully repopulated intrahepatic bile ducts in decellularized rat liver. Given the regional differences in the cell characteristics and gene profiles between extra and intrahepatic bile duct organoids [24, 25], it is reasonable to use region-specific organoids for each target of intra and extrahepatic bile ducts to establish an engineered liver graft.

For clinical applications rather than an experimental preparation of a small amount of regenerative tissue, several problems must be overcome. The immune response and the number of the cells are examples of potential issues. To avoid the immune response, an autologous cell source that does not require immunosuppression therapy is a potential solution. As liver ductal organoids can be generated from a small piece of a patient liver without gene editing, engineered bile ducts can be reconstructed using autologous cells. Moreover, the liver ductal organoids are genetically stable during long-term expansion [14], which allows the supply of a large number of cells. Because liver ductal organoids are also envisioned as a hepatocyte cell source owing to their bipotential capacity, large numbers of the organoids will be needed for transplantation as the hepatocyte cell source. To supply large numbers of cells, a highly efficient method of culturing liver ductal organoids has been reported [16]. Given these advantages, liver ductal organoids have the potential to be a cell source for bioengineered liver grafts.

We demonstrated the cholangiocyte properties of the repopulated liver ductal organoids, although functional interactions between recellularized bile ducts and hepatocytes remained ambiguous. The integrated function of the bile efflux and transportation are important in recellularization. Primary hepatocytes quickly lose their functions *in vitro* [2, 26]. Likewise, it is difficult to maintain recellularized primary hepatocytes in perfusion culture for a long time, whereas repopulated liver ductal organoids were viable for more than a week. Such a limited viability restricts the ability of recellularized hepatocytes to construct the functional structures. Thus, in future studies, it is important to improve hepatocytes as a resource for recellularization, as well as to evaluate the cellular integration between recellularized bile ducts and hepatocytes upon experimental transplantation.

In conclusion, we demonstrated that liver ductal organoids are a useful cell source for the reconstruction of intrahepatic bile ducts in a 3D scaffold of decellularized liver and to enable the appropriate cell distribution of recellularized hepatocytes and bile ducts in bioengineered liver grafts. It would be of great interest to develop liver ductal organoids as a hepatocyte source as well as to generate functional recellularized liver grafts *in vivo* for the clinical application of these cells in the future.

## 4. Materials and Methods

### 4.1. Animals and human liver samples

All the animal experiments were performed in accordance with the Animal Protection Guidelines of Kyoto University and with approval from the Animal Research Committee of Kyoto University.

Liver specimens were obtained from resected liver tissues of patients who underwent hepatectomy at Kyoto University Hospital and provided written informed consent, following the approval given by Kyoto University Graduate School and Faculty of Medicine, Ethics Committee (R1671).

### 4.2. Preparation and culture of mouse liver ductal organoids

Mouse liver ductal organoids were prepared and cultured as previously described [13, 14], but with slight modifications. Briefly, the liver was harvested from C57BL/6J mice (8–12 weeks old, female, CLEA Japan, Shizuoka, Japan), mechanically minced, and serially passed through needles for further mechanical fragmentation. The clusters were subsequently treated with 2.6 μU/mL Liberase DH (Roche, Basel, Switzerland) in DMEM (Wako, Pure Chemical Industries, Osaka, Japan) containing 0.1 mg/mL DNase I (Sigma-Aldrich, St. Louis, MO) for 10 min at 37°C. The digested fragments were serially strained using mesh filters of 100 and 40 μm (BD Falcon, Franklin Lakes, NJ, USA). The fractions on the filters were suspended in DMEM (Wako) and centrifuged at 440 × *g* for 3 min, and the pellet was embedded in Matrigel-GFR (Corning, NY).

Liver ductal organoids were cultured in Matrigel-GFR with mouse EM (Supplementary Table S1) containing 10 μM forskolin (Wako). Organoids were passaged every 4–6 days by dispersion with TrypLE Express (Invitrogen, Carlsbad, CA) [13]. The dispersed cells were re-embedded in Matrigel-GFR or used for recellularization.

### 4.3. Preparation and maintenance of human liver ductal organoids

Human liver specimens (0.5–1.0 cm^3^) were obtained from the non-tumorous part of the resected liver from patients who underwent hepatectomy. The specimens were subjected to the same procedure as the mouse organoid culture described above. For passaging, the application time of TrypLE Express was 10 min. The organoids were cultured in human EM (Supplementary Table S1) [14].

### 4.4. Isolation of mouse primary hepatocytes

Primary hepatocyte isolation was performed using two-step collagenase perfusion technique, as previously described [27]. Briefly, a C57BL6/J mouse was anesthetized with isoflurane (Wako) and the inferior vena cava was exposed and cannulated with a 25-gauge needle. After clamping the superior vena cava and cutting the hepatic portal vein, the liver was perfused with 50 mL of Ca^2+^-free HBSS containing 0.5 mM EGTA (Wako) and 2 U/mL heparin (Novo-Heparin; Mochida, Tokyo, Japan) for 7 min, followed by 100 mL of collagenase solution containing 0.2% dispase II (Sanko Junyaku, Tokyo, Japan), 0.2% collagenase type II (Gibco, Palo Alto, CA, USA), 0.1 mg/mL DNase I (from bovine pancreas, Sigma), heparin 2 U/mL, 150 mmol/L NaCl, 5.4 mmol/L KCl, 0.34 mmol/L NaHPO_4_, 0.1 mmol/L MgSO_4_, 5.0 mmol/L CaCl_2_, 4.2 mmol/L, NaHCO_3_, 5.6 mmol/L glucose, and 10 mmol/L HEPES (all of the chemical reagents other than those indicated were purchased from Wako) for 7 min at 37°C. The liver was then resected out and gently shaken to retrieve the cells. After filtering the isolated cells through a 100 μm strainer (BD Falcon, Franklin Lakes, NJ, USA), the cell suspension was washed three times by centrifugation at 50 × *g* for 3 min at 4°C. The isolated primary hepatocytes were immediately used for RNA extraction or the recellularization procedure.

### 4.5. Cell viability assay

After 3,000 single cells dispersed from liver ductal organoids were embedded in Matrigel, the organoids were cultured in EM for 5 days. Organoid viability was quantified every day from aliquot of the organoid using CellTiter-Glo® (Promega, Madison, WI) and GloMax Discover Microplate Reader (Promega).

### 4.6. Histological analysis

Organoids were retrieved from Matrigel, re-embedded in Cellmatrix Type I-A (Nitta Gelatin), and fixed in 10% formalin (Wako) overnight at room temperature. Recellularized liver grafts were fixed in 4% paraformaldehyde (Wako) for 24 h at 4°C or in 10% formalin overnight at room temperature, and then embedded in paraffin.

Paraffin-embedded sections (4 μm) were dewaxed, rehydrated, and subjected to either hematoxylin and eosin staining or immunohistochemical staining. For immunostaining, antigen retrieval was performed by autoclave for 15 min at 121°C, and the sections were by incubation in PBS containing 10% donkey serum and 0.1% Triton X-100 (Nacalai Tesque Inc., Kyoto, Japan). All antibodies were diluted with PBS with 5% donkey serum and 0.1% Triton X-100. The sections were incubated with primary antibody overnight at 4°C, followed by incubation with secondary antibody for 1 h at room temperature, and then mounted using ProLong™ Gold Antifade Mountant with DAPI (Invitrogen). The stained sections were visualized using an Olympus BX50F4 microscope (Olympus Optical, Tokyo, Japan). The antibodies used for immunostaining are listed in Supplementary Table S2.

### 4.7. Quantitative real-time PCR

Before RNA extraction, liver ductal organoids in Matrigel were collected and washed twice with PBS. To extract RNA from the recellularized liver, the recellularized liver after perfusion culture was incubated with RPMI (Wako) containing 10 mg/mL collagenase type II (Gibco) for 15 min at 37°C. The solution of dissolved liver was centrifuged, the supernatant was removed, and the pellet was processed for RNA extraction.

Total RNA was prepared from organoids and engineered liver grafts using the RNeasy Mini kit (QIAGEN, Hilden, Germany) and reverse-transcribed into cDNA using QuantiTect Rev. Transcription Kit (QIAGEN). qPCR was performed on the StepOne™ system (Applied Biosystems, Foster City, CA) using Fast SYBR® Green Master Mix (Applied Biosystems). Gene expression levels were normalized to the housekeeping gene *TATA box binding protein* (*Tbp*) for mouse cells and *ACTB* for human samples. A list of primers is presented in Supplementary Table S3.

### 4.8. Rhodamine 123 efflux assay

The rhodamine 123 efflux assay was performed as described elsewhere [23]. Briefly, organoids in Matrigel on day 3 or 4 after passage were incubated with 100 μM rhodamine 123 (Sigma-Aldrich) in HBSS (Wako) for 10 min. Then, the gel was washed with HBSS three times and incubated with EM for 30 min before imaging. To inhibit MDR1 transporter activity, the Matrigel culture was preincubated with EM containing 20 μM verapamil (Wako) for 30 min before the addition of rhodamine 123. Then, the medium was changed to HBSS containing 100 μM rhodamine 123 and 20 μM verapamil. After incubation for 5 min, the gel was washed and incubated with EM containing 20 μM verapamil. Fluorescence images of the organoids were obtained using a Leica TCS SPE confocal microscope (Leica Microsystems, Wetzlar, Germany). The fluorescence intensity of images was calculated using ImageJ (National Institutes of Health, Bethesda, Maryland, USA).

### 4.9. CFTR functional assay

The CFTR functional assay was performed based on a previous report [23]. Briefly, 1000 individual cells were embedded in a 5 μL Matrigel drop and incubated in EM for 4 days. Then, the gels were preincubated with EM containing DMSO or 60 μM Inh-172 (Cayman, Michigan, USA), a CFTR-specific inhibitor, for 3 h. The gels were subsequently incubated with EM containing 10 μM forskolin (Wako) with DMSO or Inh-172 for 24 h before imaging with a Leica DMi8 (Leica Microsystems) using the Leica Application Suite X (LAS-X) software. The swelling rate of the organoids was calculated by comparing the total area of each organoid before and after the stimulation, using ImageJ.

### 4.10. Caspase activation assay

Caspase activation was quantified after incubation for 24 h with DMSO or Inh-172, as described in *Section 4.9. CFTR functional assay*, by Caspase-Glo® 3/7 Assay (Promega) using GloMax Discover Microplate Reader (Promega).

### 4.11. Preparation of GFP-expressing organoids

pLX304-GFP, a GFP-expressing lentiviral vector, was generated by transferring a GFP coding sequence from pALB-GFP (Addgene #55759) into pENTR4 (Addgene #17424) followed by gateway cloning into pLX304 (Addgene #25890). To generate lentiviral particles, HEK293FT cells were co-transfected with pLX304-GFP, psPAX2 (Addgene #12260), and pMD2.G (Addgene #12259) using X-tremeGENE™ HD (Roche) in accordance with the manufacturer’s instructions. Viral supernatant was harvested at 48 and 72 h post-transfection and filtered through a 0.45 μm PVDF membrane (Merck Millipore). For lentiviral infection, the liver ductal organoids were dissociated into single cells and mixed with the filtered viral supernatant. The cell-lentivirus mixture was supplemented with 10 μM Y27632 (LC Laboratories, Woburn, MA), 1 mM N-acetyl-L-cysteine (Wako), and 10 μg/mL polybrene (Sigma), transferred to an untreated plate, and centrifuged at 2000 rpm for 1 h at room temperature. After centrifugation, the supernatant was discarded and the cells were embedded in Matrigel drops (5–8 μL each) and cultured with EM containing forskolin. After expansion, the organoids were passaged and selected by blasticidin (Wako).

### 4.12. Harvest, decellularization, and recellularization of rat whole liver

#### Harvest and decellularization

Harvest and decellularization of the liver were performed as previously reported [8]. Under general anesthesia with isoflurane (Wako), Lewis rats (8–24 weeks old, female, SLC, Hamamatsu, Japan) underwent laparotomy. Heparin (1 U/g of body weight) was administered intravenously. The abdominal aorta was cannulated with a 20-gauge cannula after clamping of the descending aorta in the thoracic cavity. The liver was perfused with 50 mL of PBS after the transection of the superior and inferior vena cava. The portal vein and the common bile duct were cannulated with 20-gauge and 24-gauge cannula respectively. The liver was harvested and cryopreserved at −80°C. For decellularization, the liver was thawed overnight at 4°C and then perfused with 0.25 w/v% Trypsin-1 mmol/L EDTA·4Na solution with phenol red (Wako) at 37°C for 1 h, followed by 1% polyoxyethylene(10) octylphenyl ether (Wako)/0.05% EDTA (Sigma) solution for 48 h at room temperature. The decellularized liver was sterilized by perfusion with 0.1% peracetic acid (Sigma) for 2 h and washed with sterilized PBS.

#### Recellularization

Liver ductal organoids were dissociated into single cells and 3–5 × 10^6^ cells suspended in 5 mL of EM were administered into the scaffold through the bile duct at a flow rate of 1 mL/min. Before perfusion, the recellularized liver was incubated at 37°C in EM supplemented with 50 μg/mL gentamicin (Gibco), 2.5 μg/mL of amphotericin B (Gibco), and 10 μM forskolin for 3 h. We applied our previously described recellularization protocol for primary hepatocytes [8]. In total, 5 × 10^7^ hepatocytes were suspended in 30 mL of HCM^™^ (Lonza, Sales Ltd, Basel, Switzerland) and injected via the bile duct at a flow rate of 1 mL/min. The recellularized liver was incubated at 37°C for 3 h before starting perfusion culture.

### 4.15. Perfusion culture

After incubation for 3 h following recellularization, the recellularized liver was placed in the circulation culture system, and the cannula of the portal vein was connected. The perfusion culture was conducted to continuous flow at a rate of 0.7–1.0 mL/min at 37°C.

### 4.14. 3D bile duct imaging

The liver with repopulated bile ducts of GFP-expressing cells was cultured in the circulation culture system for 5 days and imaged using confocal microscopy (Leica Microsystems). A reconstructed 3D image was prepared from Z-stack images using ImageJ (National Institutes of Health).

### 4.15. Statistical analyses

A *p* value of < 0.05 was considered to indicate statistical significance; analyzes were performed by unpaired *t*-test using GraphPad Prism 9 (GraphPad Software, San Diego, CA, USA) (Fig. 2B, D, E).

## Author contributions

Katsuhiro Tomofuji: Investigation, Data curation, Writing-original draft. Ken Fukumitsu: Conceptualization, Project administration, Methodology, Writing-review & editing, Funding acquisition. Jumpei Kondo: Methodology, Writing-original draft, Funding acquisition. Takamichi Ishii, Satoshi Ogiso, Yu Oshima: Methodology. Takashi Ito, Satoshi Wakama, Kenta Makino, Hiroshi Horie: Validation. Masahiro Inoue: Supervision, Funding acquisition. Etsuro Hatano: Supervision.

## Acknowledgements

This work was supported, in part, by a Grant-in-Aid from P-CREATE, a Japan Agency for Medical Research and Development, Japan, 19cm0106203h0004 (M.I., J.K.), by a Grant-in-Aid from Takeda Science Foundation (M.I.) and by Japan Society for the Promotion of Science (K.F.). The authors would like to thank for the technical assistance at Medical Research Support Center, Graduate school of medicine, Kyoto university. The authors would like to thank Enago (www.enago.jp) for the English language review.

## Appendix A. Supplementary data

The following is the supplementary data related to this article:

Supplementary Figure and Table.docx

Supplementary Video S1.avi

## Data availability

The raw data required to reproduce these findings are available from the corresponding authors upon reasonable request.

## Supplementary Materials

**Fig. S1.**
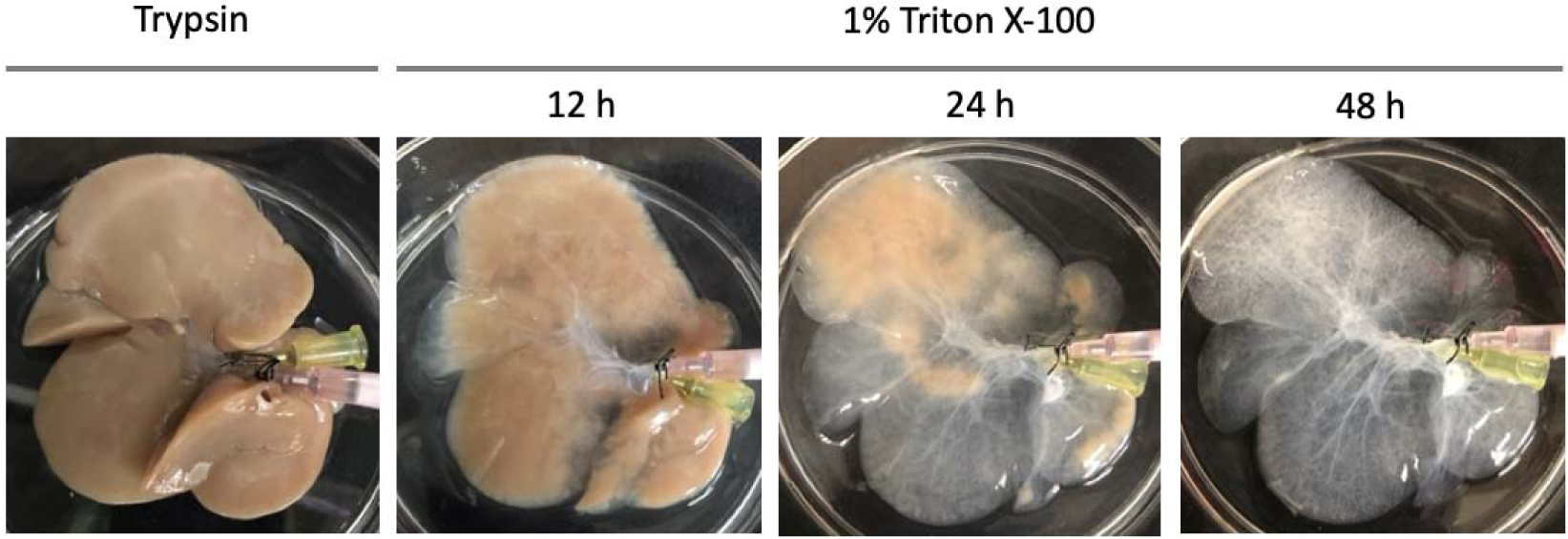
Macroscopic images of a cadaveric rat liver decellularized by portal vein perfusion.

**Fig. S2.**
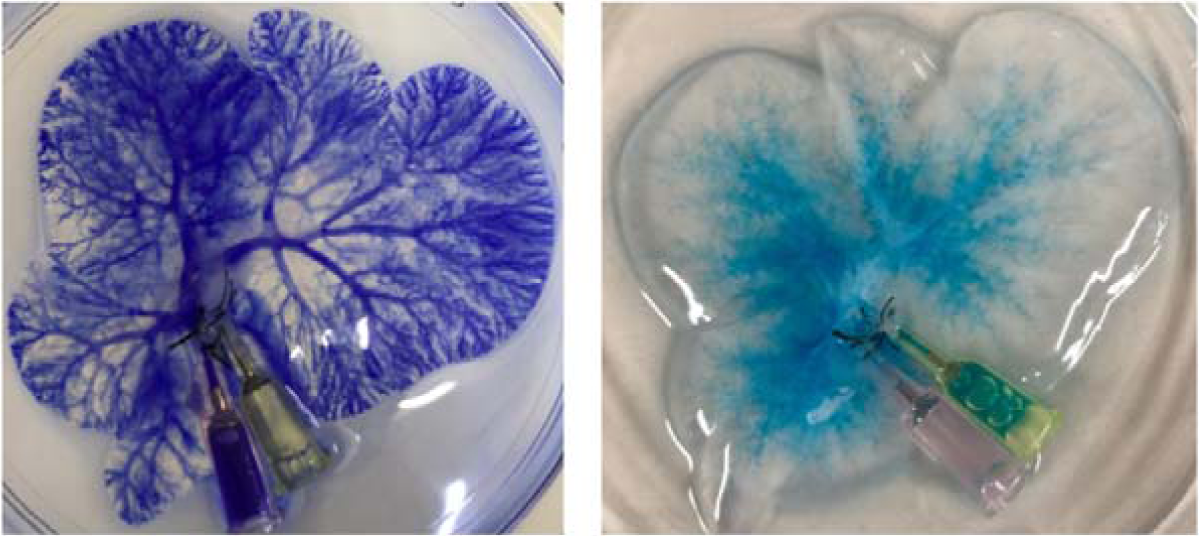
Representative photographs of intrahepatic portal veins (A) and bile ducts (B) in decellularized livers filled with crystal violet and Alcian blue solution respectively.

**Fig. S3.**
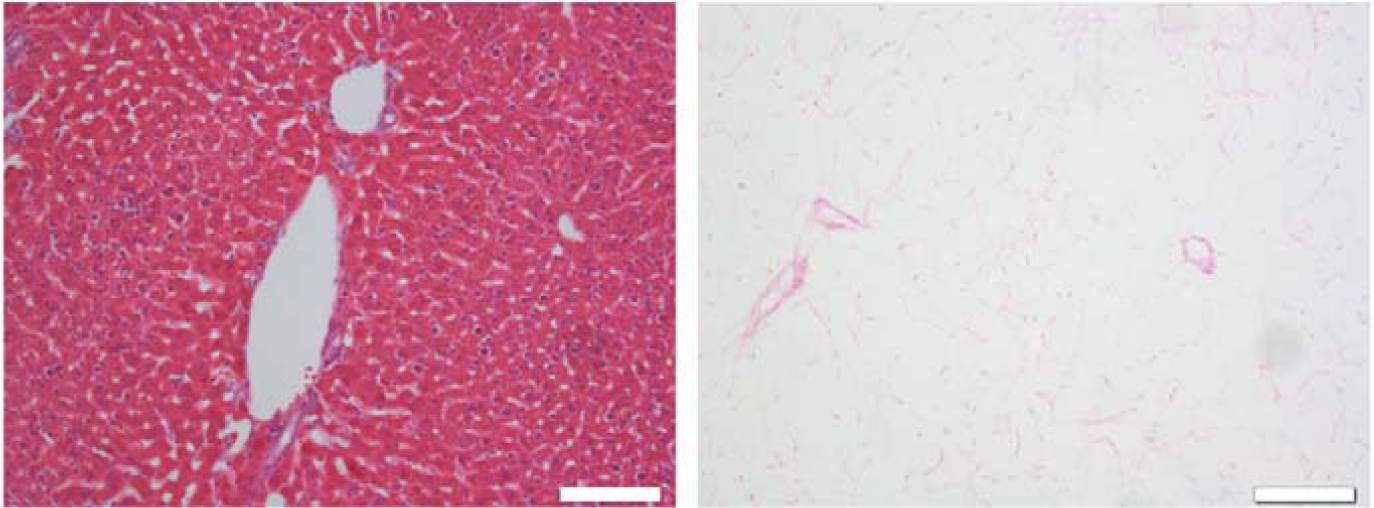
HE staining images of a native liver (A) and a decellularized liver (B). Scale bars: 100 μm.

